# *Candidatus* Rickettsia mendelii has the smallest and most ancestral-like genome in the genus *Rickettsia*

**DOI:** 10.64898/2025.12.09.692894

**Authors:** Yasuhiro Gotoh, Kentaro Kasama, Seigo Yamamoto, Shuji Yoshino, Hiroki Kawabata, Shuji Ando, Tetsuya Hayashi

## Abstract

*Rickettsia* species, which constitute obligate intracellular bacteria adapted to a host-dependent lifestyle, have traditionally been classified into four phylogenetic groups. However, the available genomes are biased toward the Typhus and Spotted Fever groups, with early-diverging Ancestral group (AG) lineages being underrepresented. Here, we isolated six *Candidatus* Rickettsia mendelii strains from *Ixodes turdus* ticks in Japan. We obtained their whole-genome sequences, including one closed genome, and found that they form a distinct lineage in the AG and have the smallest genome among the known rickettsial species. Gene tree-aware reconstruction of evolutionary events using amalgamated likelihood estimation revealed that the last common ancestor of *Rickettsia* had a smaller gene family size than modern species do, and *Ca*. R. mendelii retains the most similar genomic content to this ancestral state. Various patterns of gene gain and loss among *Rickettsia* lineages were also suggested, highlighting their divergent evolutionary trajectories. These findings increase our understanding of *Rickettsia* genome evolution.

## Introduction

*Rickettsia* species are obligate intracellular bacteria belonging to the class Alphaproteobacteria. They exhibit a biphasic life cycle involving both vertebrate and invertebrate hosts. Some species form symbiotic associations within specific arthropods, whereas others are transmitted by arthropods and cause pathogenic infections in humans. Although their classification remains under debate, *Rickettsia* species have traditionally been divided into four groups on the basis of phylogenetic analyses of several marker genes^1^: the Ancestral group (AG), the Typhus group (TG), the Transitional group (TRG), and the Spotted Fever group (SFG).

As *Rickettsia* requires host cells for cultivation and popular genetic manipulation methods are not applicable, genomic analyses have played crucial roles in elucidating their biological characteristics. The genomes of *Rickettsia* species are generally characterized by small sizes and low GC contents, reflecting adaptation to an obligate intracellular lifestyle, and are considered the products of reductive genome evolution that occurred during the transition from a free-living state to an obligate intracellular state^2^. However, the presence of mobile genetic elements and conjugative plasmids suggests that *Rickettsia* species retain the potential to acquire genes through horizontal transfer and exchange genetic material with other bacteria. Many repetitive sequences are also present throughout their genomes^3^, potentially facilitating genomic plasticity and structural rearrangements. Furthermore, comparative genomic studies have not identified clear differences in the repertoire of virulence-related factors, suggesting a possible association between the degree of genome reduction and pathogenicity^4^.

Many novel *Rickettsia* species have been reported recently, including *Candidatus* Rickettsia mendelii^5^. It was proposed as a novel species on the basis of a phylogenetic analysis of a concatenated dataset comprising the partial amino acid sequences of citrate synthetase (GltA) and the nucleotide sequences of the 16S rDNA gene. Sequences identical or closely related to *Ca*. R. mendelii have been detected in *Ixodes ricinus* (central^5–7^ and southern Europe^8^), *I. brunneus* (the United States^9^), *I. silvanus* (*Ixodes* sp. cf. *I. brunneus*) (Argentina^10^), and *I. pavlovsky* (Russian Far East^11^). These findings suggest a possible association between *Ca.* R. mendelii and tick species that infect migratory birds^9^. However, *Ca*. R. mendelii has not yet been isolated. Thus, only phylogenetic analyses based on a few gene sequences have been conducted^5, 11^. Although the analyses suggested that it belongs to the AG^11^,which occupies a basal position within the genus *Rickettsia*, genomic information for AG members remains very limited, and only those of *R. belli* and *R. canadensis* are available^12^.

Here, we isolated six *Rickettsia*-like organisms from *Ixodes turdus* collected at various locations across Japan and performed whole-genome sequencing analysis. A closed genome sequence was obtained from one of the isolates. The six isolates shared highly similar genomic features and were identified as *Ca*. R. mendelii on the basis of the sequences of *gltA* and other marker genes for *Rickettsia*. The genome of *Ca*. R. mendelii was smaller than those of TG species and *R. canadensis*, thus representing the smallest genome among the known genomes of the genus *Rickettsia*. Phylogenomic analysis based on the core genes of *Rickettsia* revealed that *Ca*. R. mendelii belongs to the AG. Furthermore, the results of evolutionary reconstruction via the use of the amalgamated likelihood estimation (ALE) method indicated that the gene content in *Ca*. R. mendelii is most similar to the ancestral state of the genus *Rickettsia* among the known genomes of this genus. Lineage-specific evolutionary trajectories within *Rickettsia* were also inferred through this analysis.

## Results

### Isolation of *Rickettsia*-like organisms and genome sequencing

Six *I. turdus* ticks were collected in five regions of Japan from 2006–2007, of which four were obtained from birds in four regions and two obtained from vegetation in Munakata, Fukuoka prefecture (Table 1). The four birds were different migratory species^13^. By inoculating tick homogenates into L929 cell cultures, we observed *Rickettsia*-like organisms and successfully obtained cell cultures of six isolates (Fig. 1A). Total DNA from these cultures was purified and subjected to whole-genome sequencing to obtain draft sequences (refer to the Methods for details). Isolate It06094 was subsequently subjected to low-input DNA-optimized transposase-based long-read sequencing to obtain a closed-genome sequence by long-read-based assembly (Supplementary Table 1). The genome of It06094 consisted of a circular chromosome of 1,050,158 bp with a GC content of 29.3% (Table 1 and Supplementary Table 1), encoding 877 protein-coding sequences (CDSs), one copy of each rRNA gene (16S, 23S, and 5S), and 33 tRNA genes. No plasmids were found. The assemblies of the other five isolates comprised 27–38 scaffolds, with total genome sizes ranging from 1,050,695 to 1,059,606 bp.

**Fig. 1:**
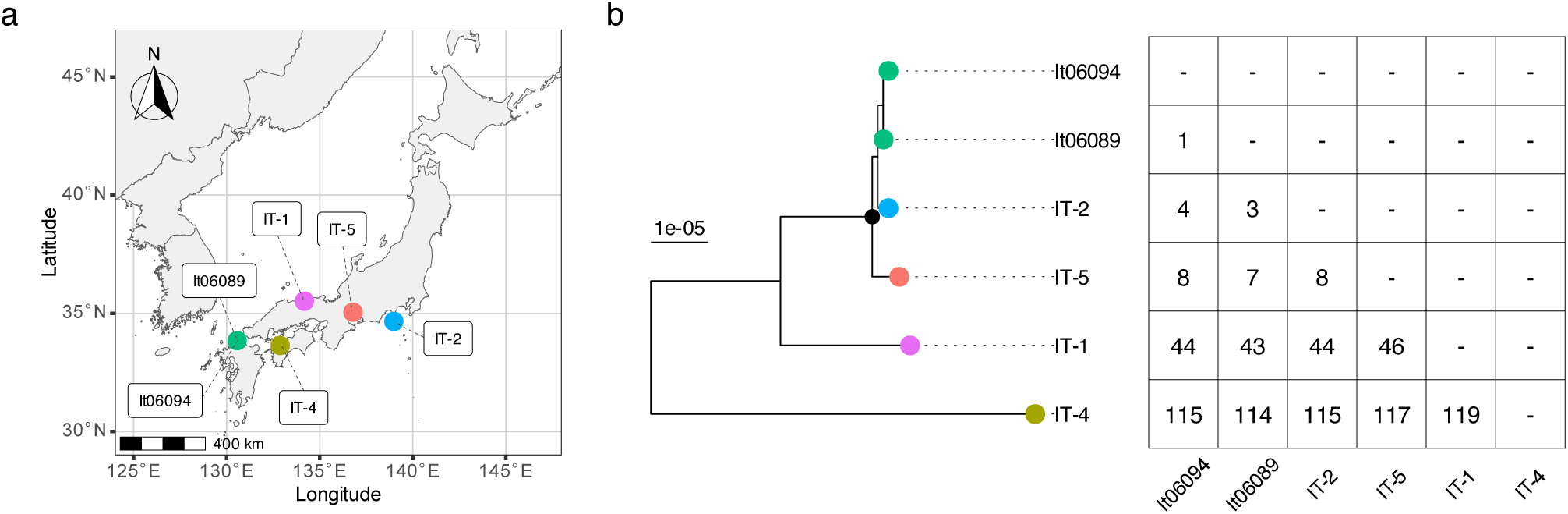
Sampling locations and genetic relationships of the six *Ca*. R. mendelii isolates. (a) Geographic information indicating the collection site of the *Ca*. R. mendelii-carrying tick. (b) Genetic relationships between the six *Ca*. R. mendelii isolates.

**Table 1:** The six *Ca*. R. mendelii isolates analyzed in this study.

| Name of Isolate | It06089 | It06094 | IT-1 | IT-2 | IT-4 | IT-5 |
| --- | --- | --- | --- | --- | --- | --- |
| Date of tick collection | April 25, 2006 | April 25, 2006 | April 4, 2007 | February 11, 2007 | November 8, 2007 | January 6, 2008 |
| Location of tick collection<br>(Latitude, Longitude) | Munakata, Fukuoka, Japan<br>(33.847, 130.567) | Munakata, Fukuoka, Japan<br>(33.847, 130.567) | Tottori, Tottori, Japan<br>(35.516, 134.176) | Shimoda, Shizuoka, Japan<br>(34.662, 138.984) | Kumakogen, Ehime, Japan<br>(33.642, 132.877) | Yatomi, Aichi, Japan<br>(35.049, 136.787) |
| Tick source | Vegetation | Vegetation | *Turdus pallidus* | *Otus lempiji* | *Emberiza spodocephala* | *Tarsiger cyanurus* |
| Tick species | *Ixodes turdus* | *Ixodes turdus* | *Ixodes turdus* | *Ixodes turdus* | *Ixodes turdus* | *Ixodes turdus* |
| Tick stage | Larva | Larva | Nymph | Larva hatched from the egg laid by the captured female tick. | Nymph | Nymph |
| Genome size (bp) | 1,051,348 | 1,050,158 | 1,056,858 | 1,050,695 | 1,059,606 | 1,057,431 |
| Number of contigs | 27 | 1 | 34 | 38 | 36 | 32 |

Self-to-self comparison analysis of the It06094 genome revealed three types of short repetitive sequences (Rep1, 985 bp; Rep2, 624 bp; and Rep3, 50 bp) (Supplementary Figure 1 and Supplementary Table 2). All repetitive sequences were found in intergenic regions (Supplementary Table 3). Rep1 and Rep2 exhibited low similarity to the IS*982* (47.3% identity) and IS*6* (41.4%) family transposases, respectively, according to a BLASTx search, suggesting that they are remnants of insertion sequences (Supplementary Table 4). Rep3 revealed no significant similarity to known sequences, and unlike the *Rickettsia* palindromic elements^3, 14^, no stable secondary structures were predicted.

### Species identification of *Rickettsia*-like organisms

To identify species, we calculated the average nucleotide identities (ANIs) between the genome sequences of the six isolates and 33 *Rickettsia* species (Fig. 2 and Supplementary Table 5). The ANI values between the six isolates ranged from 99.95 to 99.99%, much higher than the 95–96% threshold generally used for species delimitation^15^, indicating that they represent the same species. On the other hand, the ANI values between the isolates and the other *Rickettsia* species were ≤83.57%, indicating that the isolates were genetically very distant from known *Rickettsia* species; therefore, we were unable to identify the species or predict their close relatives by this analysis.

**Fig. 2:**
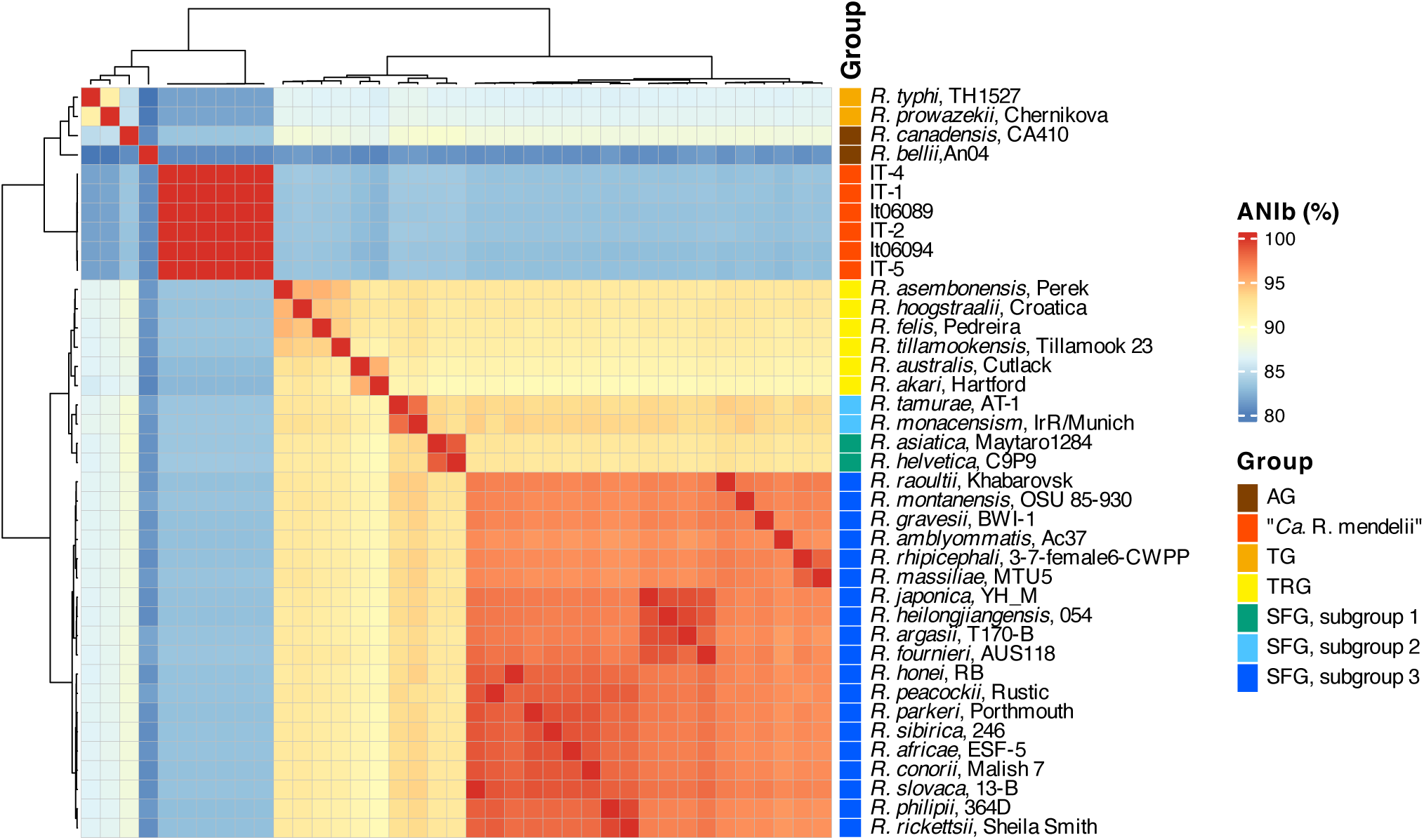
Heatmap of average nucleotide identities between 33 *Rickettsia* species and the six *Ca*. R. mendelii isolates.

Next, to include the *Rickettsia* species whose genome sequences were unavailable in the analysis, we conducted gene-level species identification using five genes frequently used for *Rickettsia* species identification [16S rRNA, *ompB*, *sca4*, *gltA* and *htrA* (also known as gene *D* or 17-kDa antigen gene); however *ompA*, which was also used for this purpose, was not identified in the six isolates, as in many non-SFG *Rickettsia* species^16, 17^]. The sequences of the five genes, showing 100% identity between the six isolates, were searched against the NCBI core_nt database using BLASTn to identify closely related species (Supplementary Table 6). The 16S rRNA, *ompB* and *gltA* sequences revealed 99.8, 99.3 and 100.0% identity, respectively, to those annotated in the description field as *Ca.* R. mendelii isolate Pr-7486_Ipav, but no record of its isolation was available, and its *sca4* and *htrA* sequences were not found in the database. Among validly published or proposed *Rickettsia* species other than *Ca*. R. mendelii, the highest sequence identities observed were 98.46% for 16S rRNA (*R. belli*), 84.7% for *ompB* (*R. aeschlimannii*), 78.7% for *sca4* (*R. slovaca*), 88.0% for *gltA* (*Ca.* R. tarasevichiae), and 88.8% for *htrA* (*R. helvetica*) (Supplementary Table 6). These results indicated that although the sequences of only a few partial gene fragments of *Ca*. R. mendelii have been reported, the isolates we obtained most likely correspond to the *Rickettsia* species proposed as *Ca*. R. mendelii. Hereafter, these isolates are referred to as *Ca.* R. mendelii.

In terms of the genetic relationships among the six isolates, isolate IT-4 was most distant from the others (114–119 single-nucleotide polymorphisms (SNPs)) (Fig. 1B). Isolates It06089 and It06094, both from the same region, differed by only one SNP, but they also showed only three SNPs to IT-2, which was obtained in the farthest region (779.5 km). Thus, no significant correlation was observed between pairwise SNP differences and geographic distance among the six isolates.

### Phylogenetic position of *Ca*. R. mendelii in the genus *Rickettsia*

After determining the root of the genus *Rickettsia* by phylogenetic analysis using 120 bacterial marker genes and *Orientia tsutsugamushi* as an outgroup (Supplementary Figure 2), we performed a phylogenetic analysis of 33 genome-available *Rickettsia* species and the six isolates on the basis of their core genes (n = 555) (Fig. 3). In the inferred phylogeny, *R. bellii* first diverged from the last *Rickettsia* common ancestor (LRCA), followed by *Ca*. R. mendelii. Another AG species, *R. canadensis*, subsequently diverged from the lineage leading to the other groups. The six isolates of *Ca*. R mendelii formed a tight cluster with a notably long branch, indicating that this species belongs to the AG and represents a lineage distinct from all known *Rickettsia* species. In the AG, the genomes of *Ca*. R mendelii (1.050–1.057 Mb) were comparable to but smaller than that of *R. canadensis* (1.150 Mb) and much smaller than that of *R. bellii* (1.58 Mb), which has the largest genome among the known *Rickettsia* species (Fig. 3 and Supplementary Table 5). The genome sizes of *Ca*. R mendelii were also comparable to but smaller than those of the TG species (1.109–1.112 Mb), which are known to have the smallest rickettsial genomes. The GC content of *Ca*. R. mendelii (29.2–29.3%) was >1.8% lower than those of *R. canadensis* (31.01%) and *R. bellii* (31.69%) and comparable to that of the TG (*R. prowazekii*, 29.01%; *R. typhi*, 28.92%) (Supplementary Table 5). These results indicated the similarity of *Ca*. R. mendelii to the TG *Rickettsia* in terms of genome size and GC content, although they belonged to distinct lineages.

**Fig. 3:**
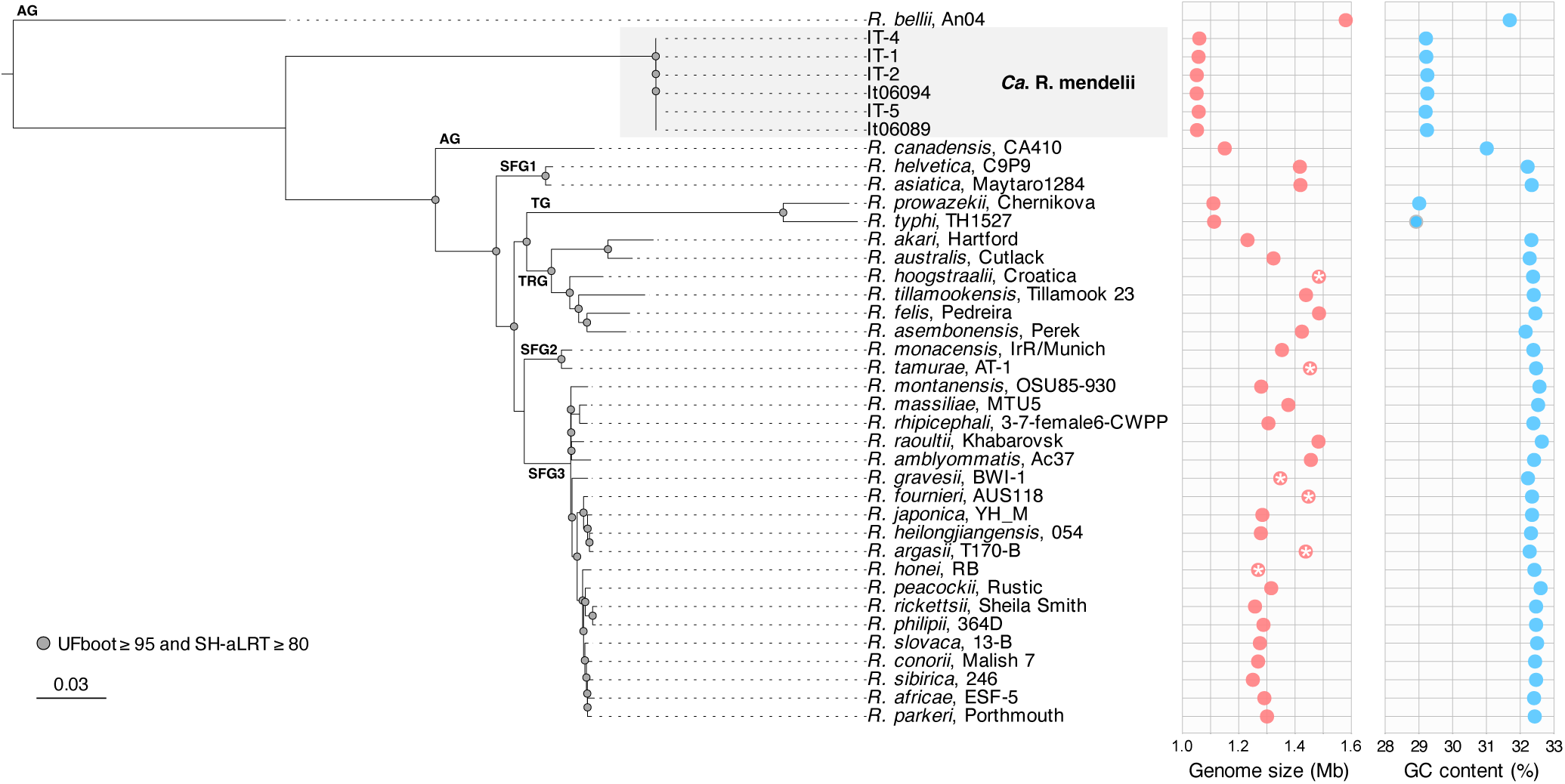
Phylogenetic position of *Ca*. R. mendelii in the core gene-based phylogenetic tree of *Rickettsia* species. A core gene-based ML tree of *Rickettsia* species was constructed. The root of the genus *Rickettsia* clade was inferred from the phylogenetic relationship between the *Rickettsia* species and *O. tsutsugamushi* (Supplementary Fig.2). Genome size (Mb) and GC content (%) are shown for each strain. The NCBI-de-fined assembly levels Scaffold and Contig are indicated by placing a white asterisk inside the circle used to plot the genome size (refer to Supplementary Table 6). Note that the clade referred to as SFGII in the reference^20^ is designated as TRG in this manuscript, and SFGI corresponds to SFG2 and SFG3 here.

In this analysis, *R. helvetica* and *R. asiatica,* which have been classified as members of the SFG^18, 19^, formed an independent clade that separated from the other SFG species before the separation of the TRG and TG (Fig. 3). These results are consistent with those provided in a report and suggest that *R. helvetica* forms an individual clade positioned basal to the TG but not the SFG ^20^. Hereafter, we designated *R. helvetica* and *R. asiatica* as SFG subgroup 1 (SFG1). In addition, among the other SFG members, *R. monacensis* and *R. tamurae* formed a clearly distinct clade and were designated SFG subgroup 2 (SFG2). The remaining SFG members were designated SFG subgroup 3 (SFG3) in this manuscript.

### Comparison of genomic structures between *Ca*. R. mendelii and other *Rickettsia* spp

*Rickettsia* genomes are characterized by a high degree of synteny conservation with a few exceptions^3, 21^. To examine the genome structure conservation in *Ca*. R. mendelii, we selected 27 *Rickettsia* strains for which closed genomes were available and compared their chromosomes with that of It06094 (Fig. 4 and Supplementary Table 5). In the AG, the genome synteny was well conserved between It06094 and *R. canadensis,* except for several inversions and translocations, whereas extensive genome rearrangements were observed between It06094 and *R. bellii*. Comparisons with other *Rickettsia* species also revealed overall synteny similar to that observed between It06094 and *R. canadensis,* except for a few cases, such as *R. peacockii*.

**Fig. 4:**
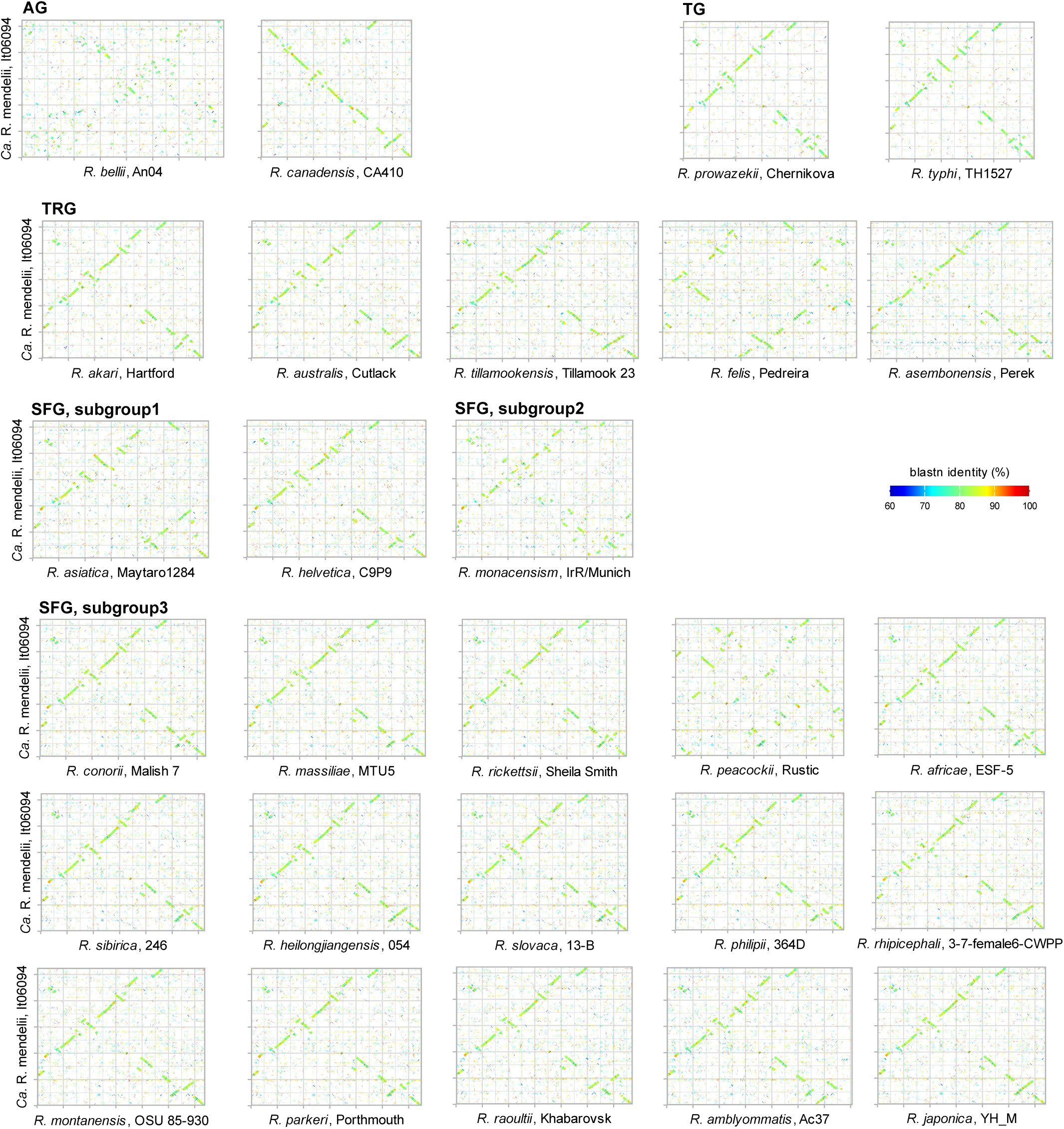
Dot plot matrix analysis of chromosome sequences between 27 fully sequenced *Rickettsia* strains and the *Ca*. R. mendelii isolate It06094. Genome sequences were rotated, when necessary, so that they begin with the *dnaA* gene on the forward strand.

### Reconstructed genome evolution process in the genus *Rickettsia*

*Ca*. R. mendelii exhibited genomic features similar to those of *R. canadensis* and the TG species but were not closely related to them in genetic or phylogenetic analyses (Figs 2 and 3). To elucidate the evolutionary processes of each lineage, we reconstructed gene family evolutionary events along the phylogenetic tree of the genus *Rickettsia* (species tree) using the gene tree-aware approach implemented in Amalgamated Likelihood Estimation (ALE)^22^. In ALE method, a sample of gene trees (e.g., bootstrap replicates) is reconciled with a given species tree to infer ancestral events (originations, duplications, intratransfers, and losses) as well as copy numbers of the gene family at all nodes of the species tree. The use of ALE increases the accuracy and robustness of gene family evolutionary inference over earlier gene tree–species tree reconciliation approaches by estimating duplication, transfer, and loss rates directly from the data while also accounting for uncertainty in individual gene trees, which are often poorly resolved. Consequently, the accuracy of ancestral reconstruction depends on the reliability of both the gene and species trees, as well as on how comprehensively the taxon sampling represents the extant diversity of the clades considered^22^.

We used the genome of It06094 together with those of 33 *Rickettsia* species to reconstruct changes in gene family content across the evolutionary history of the genus *Rickettsia* (Fig. 5 and Supplementary Table 7). We clustered 49,243 proteins from the 34 *Rickettsia* genomes into 7,103 gene families. For each family, a sample of 2,000 ultrafast bootstrap gene trees was inferred and subsequently reconciled with the species tree to reconstruct gene family histories. The ALE method outputs relative frequencies rather than integer counts for ancestral events and copy numbers. These frequencies represent statistical support, analogous to bootstrap support values for the nodes in phylogenetic trees. We adopted a relaxed minimum frequency threshold of 0.3 to capture weak but biologically meaningful signals that may be attenuated by the cumulative uncertainty arising from sequence alignment, tree inference, and reconciliation^23^. The results of the gene tree–aware ancestral reconstruction of the genus *Rickettsia* are shown in Fig. 5.

**Fig. 5:**
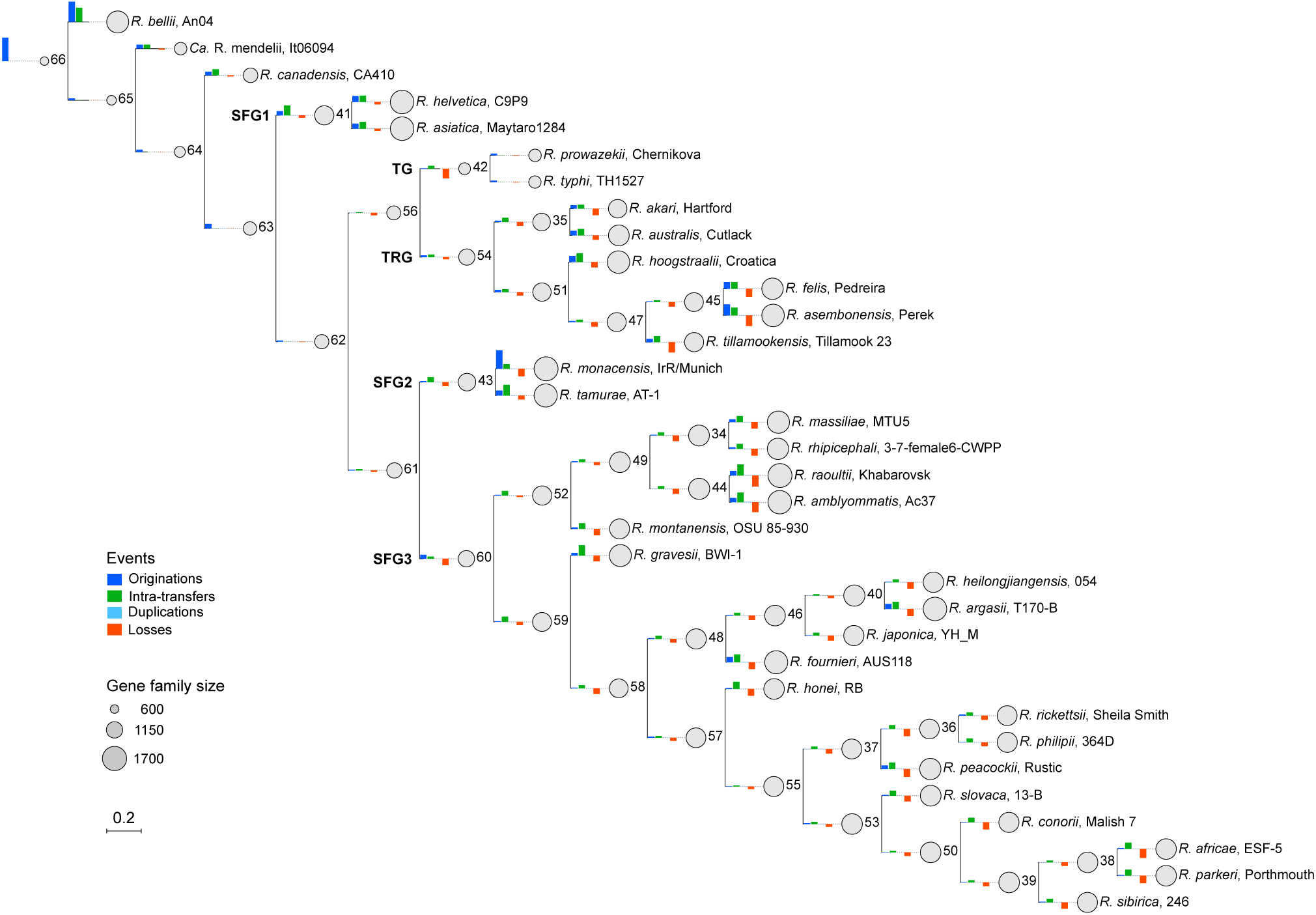
Ancestral reconstruction of gene family sizes and events in *Rickettsia* species. The phylogenetic tree includes 34 *Rickettsia* species, including *Ca*. R. mendelii It06094 and is rooted by *O. tsutsugamushi*. The sizes of gray circles at nodes and tips represent the summed gene family sizes (total inferred copy number). The heights of the bars shown along each branch indicate the numbers of originations, intratransfers, duplications, and losses. The height of the bars in the figure key corresponds to 300 events. For visualization, some branches are extended with dashed lines.

The genome of the LRCA (node 66) was inferred to contain fewer gene families (n = 597.4) than any of the 34 genomes used for this analysis. In the AG, *R. bellii* exhibited a marked increase in gene family size (n =1545), which was largely attributable to origination events (i.e., de novo genes or incoming genes from outside the species tree) and intratransfers (gene transfers within the species tree) that occurred during the evolutionary transition from the LRCA to *R. bellii*. *Ca*. R, mendelii and *R. canadensis* showed a slight increase in gene family size relative to that of the LRCA, which was driven mainly by a small number of origination and intratransfer events. However, the gene family size of *Ca*. R. mendelii” (877.0) was much closer to that of the LRCA than that of *R. canadensis* (1025.0) was, suggesting that *Ca*. R. mendelii contains a gene repertoire that is more similar to the ancestral state than *R. canadensis* does.

In the *Rickettsia* lineages other than the AG, both gene gains and losses were inferred at each node. Their gene family sizes, except for that of the TG, were notably expanded relative to that of the LRCA, while SFG3 exhibited considerable intergroup variation in gene family size [SFG1 (1549.0–1655.0), SFG2 (1623.0–1681.0), and SFG3 (1393.0–1679.0)]. In the TG, their gene family sizes were inferred to have expanded from that of the LCA of the non-AG lineage (node 63) until the TRG/TG LCA (node 56), after which a marked genome reduction occurred during the divergence from the TRG/TG LCA to the TG LCA (node 42) because of extensive gene loss. Therefore, although the TG possesses a small genome and gene family size (n = 878 in *R. prowazekii* and n = 864 in *R. typhi*), similar to *Ca*. R. mendelii and *R. canadensis*, their emergence followed an evolutionary process distinct from that of *Ca*. R. mendelii and *R. canadensis.* In the SFG1–3 and TRG lineages, both gene gains and losses occurred during the emergence of each group- or species-level LCA (internal nodes) and terminal lineage (terminal nodes). Overall, these lineages evolved toward an increase in gene family size. This trend led to larger genomes than those of *Ca*. R. mendelii, *R. canadensis*, and the TG. Their gene family sizes (n = 1297.0–1681.0) increased 2.17–2.81-fold relative to that of the LRCA.

## Discussion

In this study, we isolated *Ca*. R. mendelii from *I. turdus* and showed that it has the smallest genome among the known members of the genus *Rickettsia* and represents a distinct lineage within the AG (Fig. 3). The six isolates we obtained exhibited notably low genomic diversity, sharing more than 99.1% sequence identity with the gene sequences of *Ca*. R. mendelii available in the NCBI database. Such very low genomic diversity was also observed in *R. japonica*^24^, *R. heilongjiangensis*^25^, and *Ca*. R. kotlanii^26^, all of which belong to the SFG. The lack of correlation between SNP-based genetic distances and geographic locations among the six isolates (Fig. 1) may reflect the migratory behavior of birds parasitized by *I. turdus*^9^. *Ca*. R. mendelii has been detected in ticks collected from various regions worldwide^5, 6, 8, 9, 11, 27^. Because it has been detected in multiple *Ixodes* species, this bacterium is likely to be widely distributed through transmission mediated by migratory birds.

*Ca*. R. mendelii is presumed to be nonpathogenic^9^, as no human infections have been reported, similar to the other AG members^28^. In the genus *Rickettsia,* a paradoxical relationship between pathogenicity and genome size has repeatedly been discussed, i.e., pathogenic species often have smaller genomes than nonpathogenic species do^4, 29, 30^. However, *Ca*. R. mendelii does not follow this trend.

An interesting finding on the evolutionary processes of *Ca*. R. mendelii and other *Rickettsia* species was obtained through the reconstruction of ancestral gene content in the genus *Rickettsia* using a tree-aware approach that explicitly takes gene duplication, transfer, and loss (DTL) events into account. The results of this analysis revealed that the size of the LRCA gene family was smaller than that of any of the modern *Rickettsia* species analyzed and that the evolution of the genus *Rickettsia* has involved continuous cycles of gene gains and losses (Fig. 5 and Supplementary Table 7). Importantly, *Ca*. R. mendelii and *R. canadensis*, another AG member, experienced fewer gene gain and loss events than other *Rickettsia* lineages did. In particular, *Ca*. R. mendelii possessed the smallest genome size and gene family size among the *Rickettsia* species analyzed (Supplementary Table 7). Thus, the genome of *Ca*. R. mendelii can be considered to retain the most similar characteristics to those of the LRCA.

Although the genomes of *Rickettsia* species have traditionally been regarded as undergoing reductive evolution^2^, a recent study suggested that the genus may have followed two distinct adaptive trajectories throughout its evolutionary history^20^. *Rickettsia* genomes have evolved toward genome expansion rather than contraction, but a relatively small expansion has occurred in those of *Ca*. R. mendelii and *R. canadensis,* whereas those of the TG species underwent extensive reduction after expansion. These findings thus partly support the two trajectories insight^20^ and provide much wider views highlighting the lineage-specific evolutionary processes within the genus *Rickettsia*: Gene exchanges within and from the outside of the genus occurred frequently, but the frequencies differed notably between lineages (Fig. 5). Such differences likely arose in ecological contexts, such as mixed infections of different *Rickettsia* species within tick hosts^31^ and diverse symbiotic associations involving protozoa and other microorganisms^17^. Elucidating the biological networks that mediate these exchanges is crucial for understanding the adaptive evolution of the genus *Rickettsia*.

To better understand the evolutionary dynamics of the genus *Rickettsia* within the family Rickettsiaceae, high-quality genome sequences from newly identified *Rickettsia* lineages and from phylogenetically related species outside the genus are needed. While lineages positioned outside *R. bellii*, such as Torix spp. and *Candidatus* Megaira spp., have recently been reported^32–34^, only a few complete or near-complete genome sequences are currently available for these groups. The inclusion of these lineages in future analyses will provide much enhanced resolution in reconstructing the process in which the gene family repertoire of the LRCA was established and the subsequent evolutionary history of the genus *Rickettsia*, as well as a comprehensive understanding of the origins of pathogenicity and host adaptation in the evolutionary history of the genus *Rickettsia*.

## Methods

### *Rickettsia* detection, isolation and culture

Six *Ca*. R. mendelii isolates were obtained from *I. turdus* collected in a field investigation conducted from 2006-2007 (Table 1 and Figure 1). Ticks were collected from vegetation by flagging and from migratory birds captured for bird banding using forceps. The six isolates were obtained by inoculating tick homogenates into mouse fibroblasts (L929) as described previously^35^. All the ticks and their avian hosts were morphologically identified to species, and the tick life stages were determined under a stereomicroscope. With permission, bird captures were conducted under the following licenses issued by the Ministry of the Environment of Japan: Kankoku-Chiya-Kyo No. 070319001-38; Kankan-Chiya-Kyo No. 0612078; Kankoku-Chiya-Kyo No. 070319001-49; and Kanchu-Chiya-Kyo No. 070326001-30.

### Genome sequencing, assembly, and annotation

Genomic DNA was purified as described previously^25^. Detailed information on genome sequencing and analysis is provided in Supplementary Table 1. DNA libraries for Illumina sequencing were prepared using the following kits: (i) QIAseq FX DNA Library Kit (Qiagen) for It06089; (ii) Nextera XT DNA Library Prep Kit (Illumina) for It06094; and (iii) NEBNext Ultra II FS DNA Library Preparation Kit (New England Biolabs, MA, USA) for IT-1, IT-2, IT-4, and IT-5. These libraries were subsequently sequenced on the Illumina MiSeq platform (Illumina, CA, USA) to generate paired-end sequence reads (301 bp x2). Illumina reads were quality-filtered by trimming low-quality bases and adapter sequences using Platanus_trim (http://platanus.bio.titech.ac.jp/pltanus_trim). Illumina reads derived from mouse L929 cells were removed by mapping to the mouse reference genome GRCm39 (GCF_000001635.27) using Bowtie2 v2.4.5 ^36^ in Kneaddata v0.10.0 (https://bitbucket.org/biobakery/kneaddata). Genome assemblies were generated using Platanus_B v1.3.2^37^, which produced draft genomes of each isolate. To obtain the closed genome sequence of It06094, a long-read sequencing library was prepared using the Rapid PCR Barcoding Kit (SQK-RPB004), sequenced on an Oxford Nanopore Technologies (ONT) MinION platform using the R9.4.1 flow cell, and base-called using Guppy GPU v3.4.5 (ONT). ONT reads were mapped to the It06094 draft genome, and the aligned reads were extracted as nonmouse reads. The nonmouse reads were assembled using the microPIPE pipeline^38^, followed by short-read polishing with polypolish v0.5.0^39^. The resulting closed chromosome sequences were rotated to start at *dnaA* using Circlator v1.5.5^40^. Assembly quality assessment and gene annotation were performed using CheckM v1.2.3 ^41^ and Prokka v1.14.6 ^42^, respectively.

### Genome analyses

ANI values were calculated using PYANI v0.2.10^43^ and visualized using the ComplexHeatmap v2.22.0^44^ package in R. Genome sequence comparisons were performed via the use of BLASTn v2.13.0 ^45^ and visualized using ggplot2 v3.5.1^46^. Repetitive sequences in the It06094 chromosome were detected using REPrise v1.0.1^47^ with the parameter (-dist 2) and refined by cd-hit-est v4.8.1^48^ (parameter: -c 0.8). The chromosomal locations of the repetitive sequences in It06094 were determined using BLASTn v2.13.0 with a threshold of ≥95% identity and a ≥50 bp alignment length. Core-genome SNPs of the six isolates were identified using BactSNP v1.1.0^49^ with the It06089 finished genome as the reference. World map data were retrieved and cropped using rnaturalearth v1.0.1 in R.

### Core gene-based phylogenetic analysis

The genome sequences of *Rickettsia* species, excluding metagenome-assembled genomes, were retrieved from the NCBI RefSeq database (accessed on 9 May 2024). From the genomes containing ≤30 contigs and with completeness ≥97% and contamination ≤3%, as assessed by CheckM v1.2.3, we selected 33 genomes by applying the criterion of one representative genome per species for downstream analyses (Supplementary Table 2). These genome sequences were rotated and annotated as described for the It06089 genome. The core genes were identified using Roary v3.13.0^50^ with a threshold of 70% amino acid sequence identity. Maximum likelihood (ML) trees based on core gene sequences were constructed using IQ-TREE v2.4.0 (settings; -m MFP -B 2000 -alrt 1000 --bnni --wbtl) with a partition model^51^ and visualized using ggtree v3.10.1^52^ in R. The root position of the genus *Rickettsia* tree was inferred by a phylogenetic analysis based on the 120 bacterial marker genes identified by GTDB-tk v2.4.0^53^ using the *O. tsutsugamushi* strain Ikeda (GCF_000010205.1) as the outgroup.

### Tree-aware phylogenomic reconstruction

Of the six *Ca*. R. mendelii isolates, It06094, the genome-closed isolate, was used in this analysis. The phylogenetic relationships between It06094 and the 33 abovementioned *Rickettsia* species were inferred using the dataset of concatenated sequences of the abovementioned 555 core genes. The phylogenetic tree of the 34 *Rickettsia* species (species tree) was rooted with *O. tsutsugamushi*. (Supplementary Figure 1). Gene families comprising at least four members (n = 2,392) were identified by Roary v3.13.0, and single-gene alignments were generated using MAFFT v7.525^54^ and trimmed using trimAl v1.5.0^55^ with the -automated1 parameter. ML trees for each gene family were inferred from the trimmed alignments using IQ-TREE v2.4.0^51^ with the parameter settings of -m MFP -B 2000 -alrt 1000 --bnni --wbtl. The remaining gene families (n = 4,711) containing two or three sequences were not suitable for ML inference. However, because the ALE v1.0^22^ used for ancestral gene content reconstruction requires all gene families in a tree format, we provided mock trees for those with two or three sequences. Gene family trees were probabilistically reconciled against the species tree using the ALEobserve and ALEml_undated algorithm implemented in ALE to infer the numbers of duplications, losses, intratransfers (transfers within the sampled genomes) and originations (including both transfers from taxa outside the sampled genome set and de novo gene formation) along each branch of the species tree. Genome incompleteness was probabilistically accounted for within ALE using the genome completeness estimates obtained by CheckM v1.2.3. Singleton clusters (n = 3,640) were counted as originations for corresponding nodes. The numbers of events and gene family sizes were obtained by parsing the .uml_rec files from the ALE output using the Python package ALE_methods (https://github.com/ak-andromeda/ALE_methods). For each node, a gene family was considered present when its ‘copy’ value in the ALE output was ≥0.3. In addition, gene family evolutionary events (transfers, originations, and losses) were counted with a threshold ≥0.3. The copy numbers and evolutionary events per node or per gene family reported by ALE represent relative frequencies that reflect the probabilistic support for the occurrence and frequency of each event while incorporating uncertainty across the reconstructed gene tree samples. Gene family sizes and evolutionary events were summarized and visualized along the branches of the species tree with ETE Toolkit v4.3.0^56^.

## Data availability

The raw read data and assembled genomes obtained in this study have been deposited in the DDBJ/ENA/NCBI database under BioProject PRJDB16081. All the data supporting the findings of this study are available within the manuscript and its Supplementary Information. Additional data supporting this study are available from the corresponding author upon reasonable request.

## Acknowledgments

We thank Junya Nakamori, Toru Hara, Eiji Yamamoto, Narihito Saito, Miyako Tsurumi, Kiyoaki Ozaki and Tetsuya Ishibashi for their cooperation in collecting ticks from birds and vegetation and retrieving the collected specimens. We also thank the late Dr. Hiromi Fujita for providing *Ca*. R. mendelii isolates and Mai Horiguchi, Yuriko Sato, Toshimi Miyazaki and Mayumi Onuma for technical assistance. This work was supported by JSPS KAKENHI Grant Number JP22H04925 (PAGS) and AMED under Grant Number 22fk0108614h0202 to TH.

## Author contributions

Y.G. and T.H. conceived, designed, and supervised the study and wrote the manuscript. Y.G. and K.K. performed the experiments and data analysis. S.Ya., S.Yo., H.K., and S.A. contributed to resource provision. A.S. and T.H. contributed to funding acquisition. All the authors discussed the results and reviewed and approved the final version of the manuscript.

## Supplementary Information

### Supplementary Figures

Supplementary Figure 1: Self-to-self-dot plot matrix analysis of the genome of isolate It06094.

Supplementary Figure 2: Phylogenetic analysis of *Rickettsia* species using *O. tsutsugamushi* as the outgroup.

An ML tree was constructed on the basis of the sequences of 120 bacterial marker genes from GTDB-Tk v2.4.0 using IQ-TREE v2.4.0 (settings; -m MFP -B 2000 -alrt 1000 --bnni --wbtl).

### Supplementary Tables

Supplementary Table 1: Sequencing and assembly information of the six *Ca*. R. mendelii isolates.

Supplementary Table 2: Repetitive sequences identified in the It06094 genome (the consensus sequences are shown).

Supplementary Table 3: Locations of the repetitive sequences on the It06094 chromosome.

Supplementary Table 4: Sequences similar to those of Rep1, Rep2 and Rep3 (top 5 results from a BLASTx search against the ClusteredNR database).

Supplementary Table 5: Genome sequences of the *Rickettsia* species analyzed in this study.

Supplementary Table 6: Results of the comparison of BLASTn sequence homology against the core_nt database of five genes used for species identification.

Supplementary Table 7: ALE-inferred gene family evolutionary events and family sizes at each node.

